# CRISPR-Cas provides limited phage immunity to a prevalent gut bacterium in gnotobiotic mice

**DOI:** 10.1101/2022.05.20.492479

**Authors:** Torben Sølbeck Rasmussen, Anna Kirstine Koefoed, Ling Deng, Musemma K. Muhammed, Geneviève M. Rousseau, Witold Kot, Sabrina Sprotte, Horst Neve, Charles M.A.P. Franz, Axel Kornerup Hansen, Finn Kvist Vogensen, Sylvain Moineau, Dennis Sandris Nielsen

**Affiliations:** Section of Microbiology and Fermentation, Dept. of Food Science, Faculty of Science, University of Copenhagen, Denmark; Département de biochimie, de microbiologie et de bio-informatique, Faculté des sciences et de génie, Université Laval, Québec, Canada, G1V 0A6; Groupe de recherche en écologie buccale, Faculté de médecine dentaire, Université Laval, Québec, Canada, G1V 0A6; Section of Microbial Ecology and Biotechnology, Department of Plant and Environmental Sciences, University of Copenhagen, Frederiksberg, Denmark; Department of Microbiology and Biotechnology, Max Rubner-Institut, Kiel, Germany; Section of Experimental Animal Models, Dept. of Veterinary and Animal Sciences, University of Copenhagen, Frederiksberg, Denmark; Félix d’Hérelle Reference Center for Bacterial Viruses, Faculté de médecine dentaire, Université Laval, Québec, Canada, G1V 0A6

## Abstract

Many prokaryotes harbor the adaptive CRISPR-Cas system, which stores small nucleotide fragments from previous invasions of nucleic acids via viruses or plasmids. This molecular archive blocks further invaders carrying identical or similar nucleotide sequences. However, very few of these systems have been experimentally confirmed to be active in gut bacteria. Here, we experimentally demonstrate that the type I-C CRISPR-Cas system of the prevalent gut bacterium *Eggerthella lenta* can specifically target and cleave foreign DNA *in vitro* by using a plasmid transformation assay. We also show that the CRISPR-Cas system acquires new immunities (spacers) from the genome of a virulent *E. lenta* phage using traditional phage-assays *in vitro* but also *in vivo* using gnotobiotic (GB) mice. An increased number of spacer acquisition events were observed when *E. lenta* was exposed to a low multiplicity of infection *in vitro*, and three phage genes were found to contain protospacer hotspots. Interestingly, much less new spacer acquisitions were detected *in vivo* than *in vitro*. Longitudinal analysis of phage-bacteria interactions showed sustained coexistence in the gut of GB mice, with phage abundance being approximately one log higher than the bacteria. Our findings show that while the type I-C CRISPR-Cas system is active *in vitro* and *in vivo*, a highly virulent phage *in vitro* was still able co-exist with its bacterial host *in vivo*. Taken altogether, our results suggest that the CRISPR-Cas defense system of *E. lenta* provides only partial immunity in the gut.

## Introduction

Bacteria and archaea have the unique ability of acquiring resistance to various prokaryotic viruses (bacteriophages and archaeal viruses) and plasmids via CRISPR-Cas systems. This adaptive immunity is obtained by incorporating short fragments of DNA (spacers ∼30 nucleotides) from the invading genetic elements within the CRISPR array of the host genome (the adaptation stage). This memory of past infections enables the cell to recognize and cleave the DNA/RNA from subsequent invaders with identical or similar sequences (the interference stage) [1]. The acquired spacers are located in a clustered regularly interspaced short palindromic repeat (CRISPR) array where each spacer is flanked by direct repeats. CRISPR-associated genes (Cas) are often flanking the CRISPR arrays and are coding for proteins needed for the above-mentioned stages. The CRISPR array is transcribed and processed into CRISPR RNAs (crRNA), which will, in the interference stage, guide Cas nucleases to search and cleave nucleic acids of the invader that match the spacer, and thereby ultimately prevent infection [2–5].

During the past decade, extensive analyses of Cas proteins have revealed highly diverse CRISPR-Cas systems, which are currently classified into two large classes (Class 1 and Class 2), six types (I-VI) and numerous subtypes [6]. For example, the Class 1 type I-C CRISPR-Cas system is characterized by the following *cas* gene order *cas3-cas5-cas8c-cas7-cas4-cas1-cas2*, which are situated next to a CRISPR-array [7–10].

The microbes inhabiting the human gut (the gut microbiota, GM) play important roles in human health and disease [11]. It is therefore important to understand how bacteria defend themselves against phages in this ecosystem [12]. Most of the CRISPR-Cas research on gut-related bacteria is based on computational approaches [13–15], whereas experimental studies are sparse [16,17]. However, Soto-Perez et al. demonstrated transcription and interference activity of a type I-C CRISPR-Cas system by constructing a *Pseudomonas aeruginosa* strain carrying *Eggerthella lenta* cas genes, that subsequently was infected by *P. aeruginosa* phages [16]. A type I-C CRISPR-Cas previously found in *Bifidobacterium* spp. has recently also been described in the widespread human gut bacterium *Eggerthella lenta* [16,18]. *E. lenta* is a common member of the human GM and seems to be more abundant in individuals suffering from type-2-diabetes (T2D) [19,20] and might play a role in disease etiology via its production of imidazole propionate that impairs insulin signaling [19].

Here we investigate the functionality (adaptation and interference activity) of the type I-C CRISPR-Cas system harbored by *E. lenta* DSM 15644 against a virulent *E. lenta* siphophage PMBT5 (genome size 30,930 bp) in both *in vitro* and *in vivo* (in the gut) settings.

## Methods

### Bacterial strains, phage, and growth medium

*Eggerthella lenta* DSM 15644 (GCA_003340005.1), *E. lenta* DSM 2243^T^ (GCF_000024265.1) and phage PMBT5 (MH626557.1) were used in this study. Wilkins Chalgren Anaerobe medium (WCA, Sigma-Aldrich, St. Louis, Missouri, USA) was used for culturing as broth in Hungate tubes (Sciquip Limited, Newtown, UK), as solid media containing 1.5% (w/v) agar or as soft agar containing 0.5% (w/v) agar (Oxoid™, Thermo Fisher Scientific, Waltham, Massachusetts, USA). Anaerobic conditions were obtained as previously described [21]. Bacteria (containing cells from a single colony) were transferred from a WCA-plate to WCA-broth inside an anaerobic chamber and the bacterial cultures were subsequently incubated at 37°C for 1-3 days depending on the assay.

### Phage propagation and assays

For phage propagation, a culture of *E. lenta* DSM 15644 with an OD_600nm_ between 0.25 and 0.30 (∼ 5 × 10^8^ colony forming unites (CFU)/mL) was centrifuged for 10 min at 5 000 x g and the supernatant was discarded. The bacterial pellet was resuspended in 200 µL of 40 mM CaCl_2_ and mixed with 100 µL phage PMBT5 lysate followed by incubation for 10 min at room temperature to increase phage-adsorption. The phage-infected culture was subsequently added to either melted (52°C) WCA soft agar for plaque assay or added to WCA-broth for phage amplification. The inoculated WCA media were incubated anaerobically for 17-20 h at 37°C and OD_600nm_ was measured with Genesys™ 30 Visible spectrophotometer (Thermo Fisher Scientific).

### DNA extraction from cultures, lysates, and feces

DNA extraction from bacterial cultures, phage lysates, and fecal pellets were performed using the Bead-Beat Micro AX gravity kit (A&A Biotechnology, Gdańsk, Poland) following the protocol of the manufacturer. Purified DNA was stored at -80°C. A negative control representing *E. lenta* DSM 15644 with its native spacers along with a contamination control, consisting of autoclaved MilliQ water (Millipore corporation, Burlington, Massachusetts, USA), was included throughout all DNA extractions and PCR-steps.

### Primer design

Primers were designed with Geneious Prime v. 2019.0.4 and motif search was performed to ensure unique primer binding sites on the genome of *E. lenta* DSM 15644. Primer specificity was tested *in silico* using NCBI primer-BLAST with strict parameters as described previously [22]. Primers (Thermo Fisher Scientific) are listed in Table S1.

### CRISPR-Cas interference assay

An interference assay was designed using plasmid pNZ123 [23] containing two different spacers originating from the native CRISPR array of *E. lenta* DSM 15644 (Spacer 2 (S2): 5’- TCAGATTGTCGGGGTTGCCTGTCCCGCCTATCG-3’, Spacer 1 (S1): 5’- AATCGAATCTTCGCCCTTGCGGCCGAAAACCGG-3’) which were flanked by different protospacer adjacent motifs (PAMs) (Table S1). Two native spacers were included in the construct to increase the interference activity. Based on literature investigating type I-C CRISPR-Cas systems [16,24,25], we tested the interference activity with two different PAMs (5’-GGG, and 5’-TTC), since no experimental data identifying a functional PAM for the type I-C CRISPR-Cas system in *E. lenta* DSM 15644 were available at that time. The pNZ123-derivatives were generated with the Gibson Assembly® Cloning kit (NEB, Ipswich, Massachusetts, USA) and thereafter transformed into *E. lenta* DSM 15644 by electroporation as previously described [26]. Plasmid constructs were confirmed with PCR and Sanger sequencing (Macrogen, Amsterdam, Netherlands). A minimum inhibitory concentration test (Table S2) showed that *E. lenta* DSM 15644 was sensitive to chloramphenicol (Sigma-Aldrich) at a concentration above 1 µg/mL, thus 5 µg/mL chloramphenicol was used in media to select cells that were transformed with a plasmid (pNZ123) carrying a chloramphenicol-resistance gene [23].

### Detection of spacer acquisition

“CRISPR adaptation PCR technique using reamplification and electrophoresis” (CAPTURE) was applied to detect expanded CRISPR arrays with increased sensitivity [27] in the type I-C CRISPR-Cas system harbored by *E. lenta* DSM 15644. The CAPTURE protocol is based on an initial PCR amplification followed by a reamplification (nested PCR) with primer sets representing different strategies (internal, degenerate, repeat) [27]. PCRs were performed on SureCycler8800 (Agilent Technologies, Santa Clara, California, USA) following the CAPTURE protocol [27] using DreamTaq Green PCR Master Mix (Thermo Fisher Scientific), but annealing temperatures were adjusted to fit the designed primer sets (Table S1). After the initial PCR amplification, the PCR products were migrated on an 2% agarose gel suspended in 0.5X TBE buffer (45 mM Tris-Borate, 1 mM EDTA) at 110 Volts. The 1-kb plus DNA ladder (Thermo Fisher Scientific) was used as marker. Only every second lane was loaded with sample to minimize between-sample contamination. A sterile scalpel was used to cut out a fraction of the gel, with no visible band, that represented PCR-products with a DNA size ranging from 200-400 bp. The expected size for a single spacer acquisition in *E. lenta* DSM 15644 was 254 bp for the initial PCR (Table S3). The PCR-products were thereafter extracted from the gel with GeneJet Gel Extraction kit (Thermo Fisher Scientific) as recommended [27]. Reamplification was performed with the degenerate primer set according to the CAPTURE protocol [27] (Figure S1). In a volume ratio of 2:1, AMPure XP bindings beads (Beckman Coulter, Brea, California, USA) were used to clean the extracted PCR products to remove DNA fragments (< 100 bp) before library preparation.

### Gnotobiotic mice study

Twelve gnotobiotic (GB) outbred Swiss-Webster mice (Tac:SW, Taconic Biosciences A/S, Lille Skensved, Denmark) were bred at Section of Experimental Animal Models (University of Copenhagen) in an isolator and represented 8 female and 4 male animals. They were divided into 3 groups of 4 and housed two-by-two according to the same sex (Table S4): *E. lenta* (EL) + PMBT5 (EL+Phage, n=4), *E. lenta* + SM buffer (100 mM NaCl, 8 mM MgSO_4_, 50 mM Tris-Cl) (EL+Saline, n=4), and a baseline (as GB control, n= 4) that were sacrificed at age 3 weeks (Figure 1). The remaining 8 mice were transferred to the Department of Experimental Medicine (University of Copenhagen) in individual ventilated cages at age 5 weeks. Cage and housing conditions were as previously described [28]. The cages were sterilized and mounted to a sterile ventilation system. Animals were provided sterilized water and *ad libitum* low-fat diet (LF, D12450J, Research Diets, New Brunswick, New Jersey, USA). After two weeks of acclimatization (i.e. 7 weeks of age), the mice were ear-tagged, weighed, and individual feces were sampled. Next, the EL+Phage mice were orally administered with a mixture of bacterial host-phage cultures (*E. lenta* DSM 15644 and PMBT5) at a MOI of 1 (total 3×10^7^ CFU and PFU). With a volume of 40 µL, the bacteria and phages/saline were mixed in the ratio of 1:1 before being deposited on the tongue of the mice. This procedure was repeated after 6 h for a second inoculation. The bacterial cultures were in their exponential phase when orally administered to the mice and grown anaerobically prior to inoculation. Individual feces were then sampled (Figure 1a) along with body weight measurements (Figure S2) until the end of the experiment. Mouse feces were sampled at day 1 (before first inoculation), day 1.5 (6 hours after first culture inoculation), day 2, day 3, day 4, day 5, day 12, day 19, and day 26. As controls, feces were also sampled when transferred from isolator to individual ventilated cages (arrival) and from baseline mice prior euthanization. All samples were stored at - 80°C. The mice were euthanized by cervical dislocation at 10 weeks of age after anesthesia with a mixture of hypnorm (Apotek, Skanderborg, Denmark) and midazolam (Braun, Kronberg im Taunus, Germany) as described earlier [28]. Handling of mice during sampling were performed aseptically with the disinfectant VirkonS® (Pharmaxim, Helsingborg, Sweden) as recommended by the manufacturer. The germ-free status was initially evaluated by the size of the cecum (enlarged) of the baseline mice and culture plating (no growth) confirming the germ-free status of the mice. We also performed qPCR with universal primers targeting the 16S rRNA gene and sequenced the full 16S rRNA gene profile of fecal samples obtained at selected time points during the study. Of note, also before inoculation of the germ-free mice qPCR and 16S rRNA gene sequencing showed signs of bacterial contaminants (Figure S3). Given the enlarged cecum and absence of growth by plating we speculated that these signs reflect dead bacteria killed by sterilization of the feed. Procedures were carried out in accordance with the Directive 2010/63/EU and the Danish Animal Experimentation Act (license-ID: 2017-15-0201-01262).

**Figure 1:**
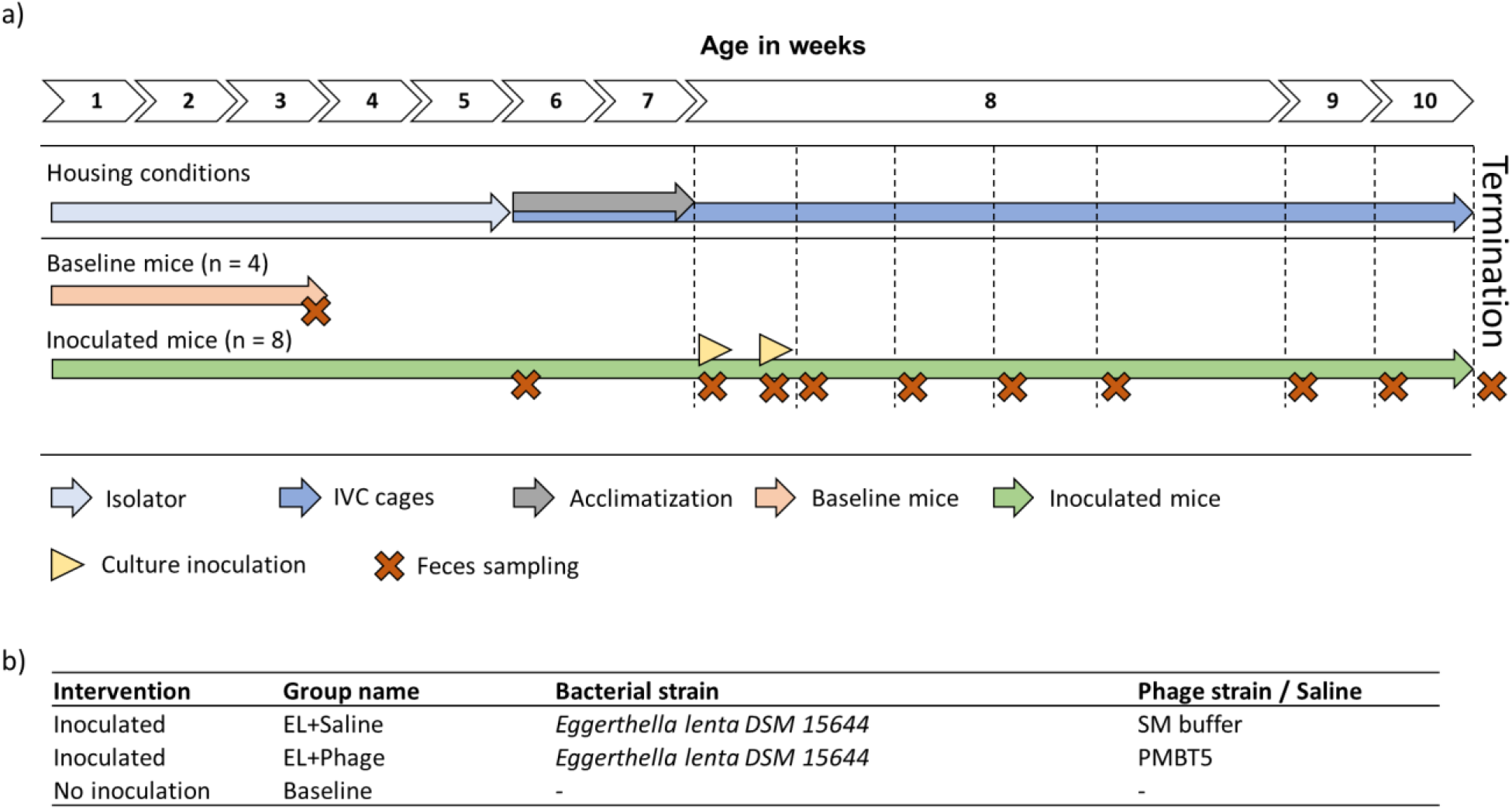
Timeline of the gnotobiotic mouse model. a) Showing the lifespan of the mice included in the study. The mice were initially bred and housed in a germ-free isolator (light blue arrow) until age of 5 weeks when they were transferred to IVCs (dark blue arrow) for individual group caging followed by two weeks of acclimatization (grey arrow) prior intervention at age 7 weeks. Feces (brown cross) were sampled from each individual mouse before and after inoculation (yellow triangle) with phages and/or bacteria. The baseline mice were euthanized and sampled at age of 3 weeks. b) Listing of the experimental groups, their abbreviation, and the inoculated bacterium and/or phage. SM buffer was used as saline solution.

### Sequencing of PCR-products

Sequencing was performed with Illumina NextSeq 550 using v2 MID output 2×150 cycles chemistry and barcodes as earlier described [29]. Illumina adaptors were designed specifically for *E. lenta* DSM 15644 (Table S1). To ensure the quality of the samples, additional cleaning with AMPure XP binding beads (Beckman Coulter), assessment of PCR-products size by gel electrophoresis, and DNA concentration measurements with Qubit HS® (Thermo Fisher Scientific) were performed between each PCR step prior to Illumina sequencing. The average sequencing depth was 231 637 reads (minimum 54 123 reads and maximum 340 311 reads) for the *in vitro* samples, and 112 927 reads (minimum 15 138 reads and maximum 320 818 reads) for the *in vivo* samples (Accession: PRJEB47947, available at ENA). Full 16S rRNA gene sequencing was performed with the MinION platform from Oxford Nanopore Technologies (ONT, Oxford, UK), as previously described [30].

### Processing of raw sequencing data

Paired ends of raw sequencing reads were merged with Usearch 11.0.667 [31] (-fastq_mergepairs) with default settings to ensure overlapping sequences of the forward and reverse reads. Subsequently, redundant sequences of primers and Illumina adaptors were removed with cutadapt 2.6 [32] (Figure S3).

### Bioinformatic analysis of sequencing and genomic data

The alignment package BWA [33], which is based on Burrow-Wheeler transformation, was used for alignment of short Illumina reads against the phage PMBT5 genome and visually interpreted with the use of Tablet 1.21.02.08 [34]. Samples with ≤ 30 reads that could be assigned to the PMBT5 phage genome were not considered, due to the numerous PCR cycles [27] and the cut of gel fragments that might have introduced minor contaminations. Local BLASTn [35] was used to match spacers originating from the type I-C CRISPR array of *E. lenta* DSM 15644 to viral genomes in the HuVirDB [16]. WebLogo [36] was used to visualize PAM sequences. CRISPRDetect [37] was used to identify CRISPR-Cas systems in genome sequences. The database of potential anti-CRISPR (acr) protein [38] was used to screen for acr proteins encoded by phage PMBT5 by the “-pblast” option with default settings in the alignment tool DIAMOND [39], and visualized in CLC Sequence viewer 8.0. The requirements of potential acr protein candidates were set to a minimum 40% of the amino acid (AA) identity sequence, length at minimum 100 AA, and for the alignment to contain shared domains with contiguous sequences.

### High-throughput qPCR (HT-qPCR) assays

The BioMark HD system was used for qPCR analysis with a Flex Six IFC chip (Fluidigm, San Francisco, California, USA) as previously described [22]. For bacteria and phage quantification, strain specific primers (Table S1) were designed to target the *cas1c* gene (NCBI GeneID: 69511386) in *E. lenta* DSM 15644 and a putative tail encoding gene (NCBI GeneID: 54998184) in PMBT5. A universal 16S rRNA primes targeting the V3-region was used as a control (Table S1). The quality of the primers was evaluated with AriaMX Real-time and Brilliant III Ultra-Fast SYBR® Green Low ROX qPCR Master Mix (Agilent Technologies) prior HT qPCR analysis as earlier described [22]. Bacterial culture of *E. lenta* DSM 15644 (5×10^8^ CFU/mL – OD_600nm_ = 0.27) was mixed with feces from germ-free mice prior DNA extraction to ensure that the genomic DNA used for the standard curve was treated as the investigated samples. The criteria for including a primer set for qPCR analysis was absence of primer dimers, no additional PCR fragments (evaluated by the melting curve), and a standard curve with efficiency between 98-102%, R^2^ > 0.991, slope ∼ -3.2, and intercept around 38. Samples with less than 10 gene copies were discarded from the analysis.

## Results

In this study we investigated the activity of the type I-C CRISPR-Cas system (Figure 2) harbored by *Eggerthella lenta* DSM 15644, when the bacterial cells were infected with the virulent phage PMBT5 during either *in vitro* or *in vivo* settings. To investigate if the type I-C CRISPR-Cas system has previously acquired spacers from other phages, we aligned the 25 native spacers in the CRISPR array with the HuVirDB (Human virome database) [16]. Only three spacers (S18, S9, and S7) were assigned to 7 viral contigs in the HuVirDB (Table S5), which was further supported by the spacers matching two recently assembled phage genomes [40]; S18 matched a *Siphoviridae* isolate (GenBank ID: BK046045.1) and S9 and S7 an unknown phage (GenBank ID: BK052885.1) [40]. None of the native spacers of *E. lenta* DSM 15644 matched phage PMBT5 genome (Table S6).

**Figure 2:**
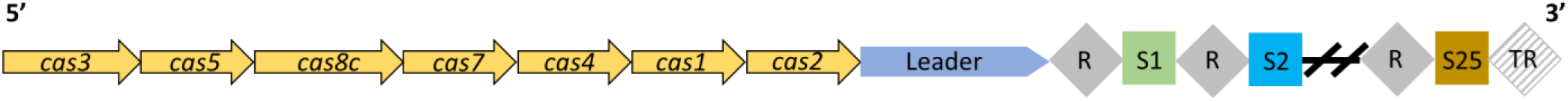
The order and structure of the type I-C CRISPR-Cas system found in *E. lenta* DSM 15644. R = Repeat, S = Spacer, TR = Terminal repeat.

### Type I-C CRISPR-Cas system of *E. lenta* can acquire new immunities *in vitro* and the new spacers preferentially target three hotspots of phage PMBT5 genome

The infection of *E. lenta* DSM 15644 with the virulent phage PMBT5 was assayed at four different MOI for 144 hours (Figure 3a). The bacterial cultures infected at MOI 10 and 1 grew to a significantly (t-test, p<0.005) higher cell density (OD_600nm_ = 0.16-0.17, approx. 48 hours after infection) compared to MOI 0.1 and 0.01 (OD_600nm_ = 0.04-0.07) (Figure 3). The cell density of the phage-infected cultures at MOI 10 and 1 was still markedly decreased (t-test, p<0.005) compared to the bacterial cultures with no phages. Different *in vitro* assays (Supplemental Methods) were performed to try to isolate CRISPR-protected bacteriophage insensitive mutants as well as plasmid interfering mutants, but to no avail (Figure S4). However, deep sequencing of PCR-amplified CRISPR-arrays from *E. lenta* DSM 15644 revealed 13 newly acquired spacers that matched phage PMBT5 genome in cultures with all four MOIs (Figures 3 and S5). The size of the acquired phage-associated spacers varied from 29-37 bp. The matching 13 unique protospacers are located in the genes coding for a phage terminase, portal-, minor structural-, adsorption-, or several uncharacterized proteins (Figures 3c and 3d, & Table S7). Based on these 13 protospacers, the PAM was predicted as 5’-TTC, but no clear motifs could be detected in the flanking sequences on both sides of the protospacer (Figure S6, & Table S7). Interestingly, 3 out of the 13 phage protospacers appeared as hotspots since they together represented 91.7% of the reads (174 637 reads out of 190 317 reads) matching the phage genome in all four MOIs (Figures 3c and 3d). These three protospacer hotspots were found within the coding sequences of a portal protein and two hypothetical proteins (YP_009807283.1, YP_009807291.1, and YP_009807318.1, Table S7). The ratio of spacer acquisitions from the hotspots varied notably between the MOIs, e.g. the fraction of spacer acquisitions targeting a hypothetical protein (YP_009807291.1**) was 11.5%, 92.6%, 92.6%, and 14.4% for MOI 10, 1, 0.1, and 0.01, respectively (Table S7).

**Figure 3:**
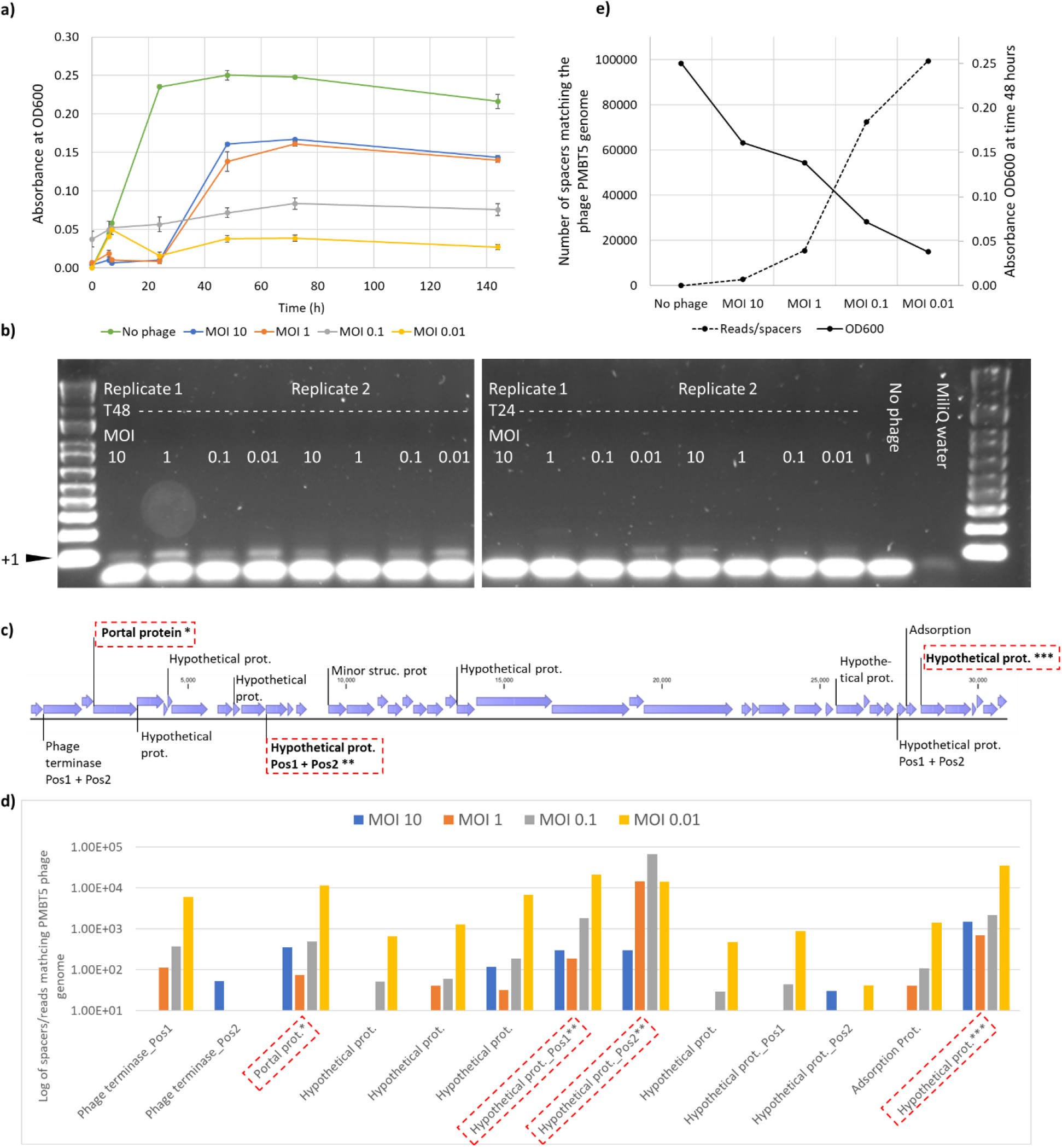
Overview of spacer acquisitions in the *in vitro* settings. a) Growth curve of *E. lenta* DSM 15644 during infection with phage PMBT5 at four different multiplicities of infections (MOI) and a control with no phages added. Bacterial growth was measured at several time points (absorbance at OD_600nm_) for 144 hours. b) Expanded CRISPR arrays in selected samples (Figure S8 for all samples) representing two replicates of all four MOI after 48 hours and 24 hours of incubation of *E. lenta* DSM 15644 exposed to phage PMBT5. DNA ladder is a 100-bp scale. With the degenerate primers, the expanded CRISPR array with one spacer “+1” was expected to yield a PCR product at ∼110 bp (Figure S1). No expanded CRISPR arrays were observed in samples with no added phages (after 48 hours incubation) or with MilliQ water added. The PCR-product at ∼40 bp likely represented primer dimers. c) The annotated genome of phage PMBT5 highlights the genes that are presented in d) with a bar plot showing the number of reads/spacers that matched to phage genes at MOI 10, 1, 0.01, and 0.01. Three genes appeared as hotspots of spacer acquisitions (coding for the portal protein (YP_009807283.1*) and two hypothetical proteins (YP_009807291.1** and YP_009807318.1***) and are marked by boxes with red dashed lines. A few genes were targeted at different positions (Pos) within the same gene. e) Graph illustrating a tendency of an inverse relation between MOI and cell density (OD_600nm_) of reads/spacer acquisitions in *E. lenta* DSM 15644 exposed to phage PMBT5.

Only a low number of reads matched the phage PMBT5 genome at MOI 10 and 1 (MOI 10: 2 648 reads (1.4%), MOI 1: 15 832 reads (8.3%) of total 190 317 reads), suggesting that at these MOIs the predominant phage resistance mechanism in these cultures was not related to CRISPR-Cas immunity. The bacterial cultures infected at MOI 0.1 and 0.01 grew only to a low cell density, yet a relatively high fraction of persisting cells acquired new spacers that matched the genome of phage PMBT5 (MOI 0.1: 72 397 reads (38.0%), MOI 0.01: 99 440 reads (52.2%) of total 190 317 reads). Interestingly, the number of spacer acquisitions matching the phage PMBT5 genome was almost linear from MOI 10 to 0.01, while bacterial biomass as determined by OD_600nm_ had an inverse tendency with a decreased growth from MOI 10 to 0.01 (Figure 3). This suggested that a low MOI may favor the adaptation activity of the type I-C CRISPR-Cas system. Taken altogether, the type I-C CRISPR-Cas system of *E. lenta* DSM 15644 can acquire new spacers from an invading phage genome.

Sequencing of all samples yielded a total of 12 276 803 reads of which 1.55% (190 317 reads) contained spacer acquisitions events that could be assigned to phage PMBT5 genome, but only one new spacer acquisition at the time. The remaining reads (98.45%) could be assigned to PCR products with no spacer acquisitions (primer dimers, 74%) and chromosomal DNA from *E. lenta* DSM 15644 (24.45%). The reads assigned to the chromosomal DNA covered the native CRISPR array (positions 1 572 740 to 1 574 444 bp) and showed a 100% nucleotide identity to 24 out of 25 spacers (Figure S7). No reads matched other parts of the bacterial DNA. This phenomenon was observed at all four MOIs as well as with the control without phage, suggesting that it did not dependent on the presence of phages.

### Efficient interference activity of the type I-C CRISPR-Cas system

A plasmid interference assay was also conducted to further evaluate the functionality of the type I-C CRISPR-Cas system of *E. lenta* DSM 15644. Two protospacers, matching S1 and S2 from the native CRISPR-array of *E. lenta*, were cloned into the vector pNZ123 with one of two PAMs (TTC or GGG) and introduced into *E. lenta*. This yielded three transformants (15644-pNZ123::GGG-S2- GGG-S1, 15644-pNZ123::TTC-S2-TTC-S1, 15644-pNZ123::WT). While the 5’-TTC motif was identified in our above phage assays, other studies [16,24,25] had suggested that GGG may be the PAM of the type I-C CRISPR-Cas system of *E. lenta*. Note that plasmid pNZ123 provides chloramphenicol resistance to the bacterial transformants. If the interference complexes of the type I-C CRISPR-Cas system recognize and cleave the two protospacers (S2 and S1), cloned into the pNZ123 vector, the chloramphenicol resistance will be lost and these bacterial transformants will be sensitive to the antibiotic. The efficiency of transformation (CFU/µg DNA) was clearly reduced (> 5 logs) with the two recombinant plasmids pNZ123::GGG-S2-GGG-S1 and 15644 pNZ123::TTC-S2-TTC-S1 compared to the control pNZ123::WT (Figure 4). These data indicate that the type I-C CRISPR-Cas system of *E. lenta* is also functional against plasmid invasion and has PAM flexibility.

**Figure 4:**
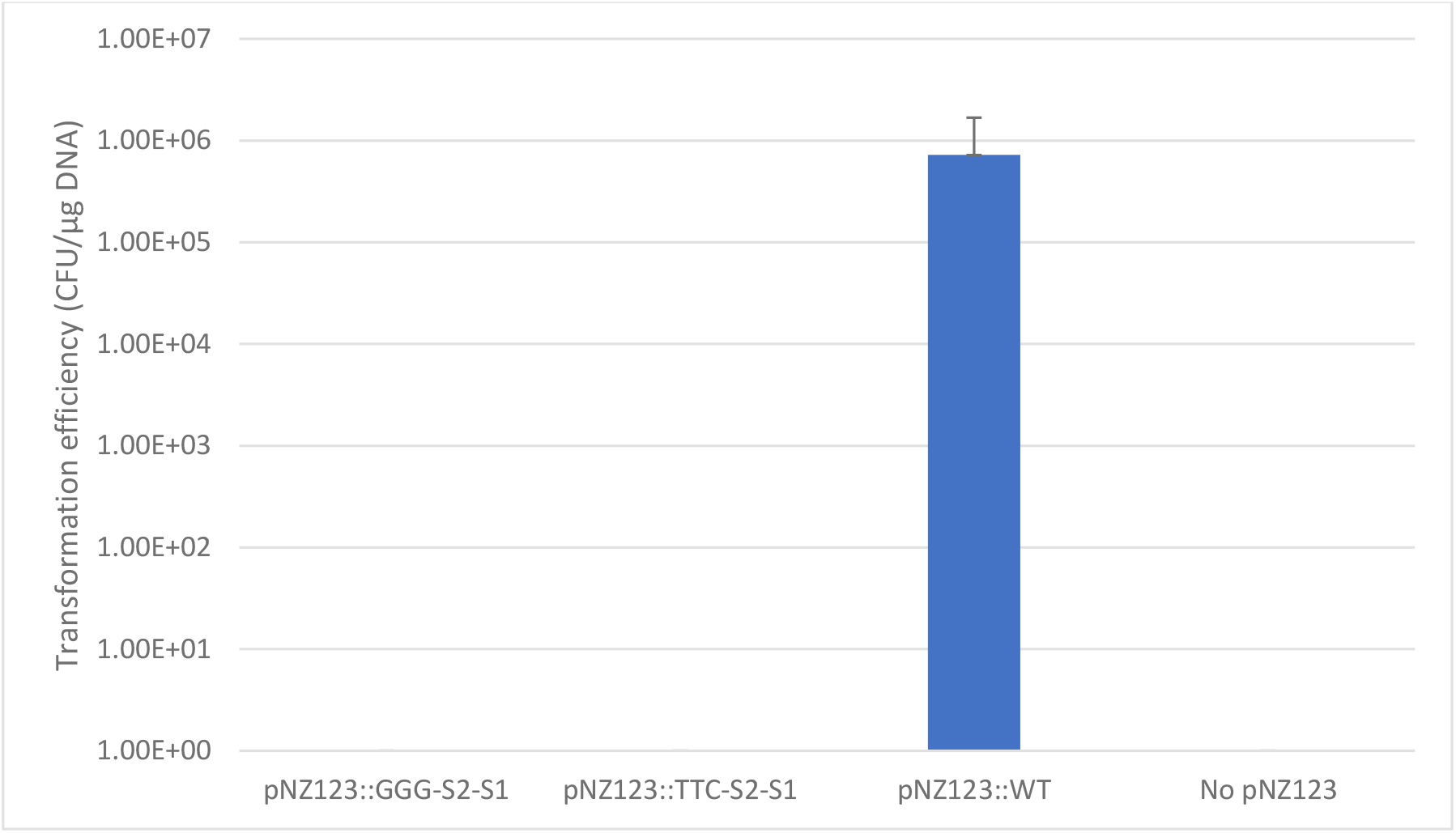
Bar plot showing colony forming units per µg DNA (CFU/µg DNA) in a logarithmic scale of transformed *E. lenta* DSM 15644 cells with plasmid pNZ123 and derivatives that provides chloramphenicol resistance. *E. lenta* DSM 15644 was transformed with pNZ123 (WT) and two derivatives containing each the same two protospacers but a different PAM (pNZ123::GGG-S2-GGG-S1, pNZ123::TTC-S2-TTC-S1, pNZ123::WT). Absence of plasmid transformation indicates interference activity of the type I-C CRISPR-Cas system. Transformation assays were performed 2, 2, 4, and 4 times, respectively.

### Co-existence of *E. lenta* and phages in the gut of gnotobiotic mice

While we could see spacer acquisition events *in vitro* when *E. lenta* DSM 15644 was infected with phage PMBT5, this study also aimed to explore CRISPR-Cas activities *in vivo*. In total, 12 gnotobiotic (GB) mice were used to (i) investigate the coexistence of *E. lenta* DSM 15644 and phage PMBT5, and to (ii) see if the type I-C CRISPR-Cas system contribute to phage resistance. The mice received either a mixture of *E. lenta* (EL) and phages (EL+Phage) or EL and saline (EL+Saline) (Figure 1). The abundance of bacteria and phages was estimated by qPCR with specific primers (DSM15644-Cas1, PMBT5-Tail). EL+Phage mice showed sustained co-existence of bacteria and phages throughout the study (Figure 5), however, at day 26 both simultaneously decreased in abundance. Phages appeared consistently to be 1 log higher compared to its bacterial host until day 19. Interestingly, *E. lenta* could co-exist with its antagonist virulent phage, since the bacterial abundance detected in the EL+Phage were comparable to the EL+Saline mice (Figure 5).

**Figure 5:**
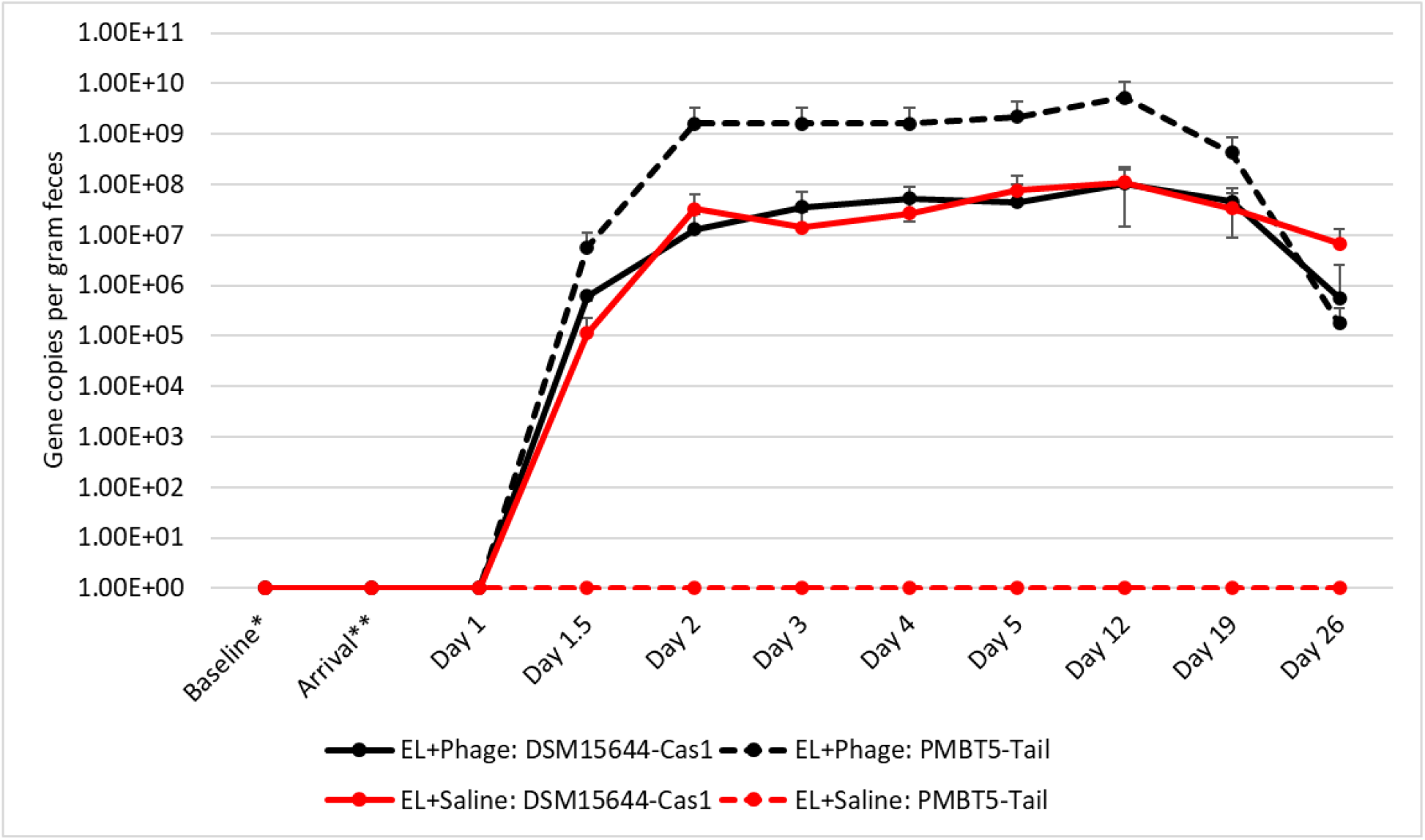
The bacterial and phage abundance in feces samples at different time points and measured by qPCR. Primers designed to specifically target the genomes of *E. lenta* DSM 15644 (*cas1* gene) and phage PMBT5 (gene coding for a putative tail protein) were used to measure total gene copies found in the feces samples. A minimum threshold of 10 gene copies was applied. *Feces samples from GB mice euthanized at the age of 3 weeks, ** feces samples from GB mice when transferred from isolator to individual ventilated cages at another housing facility.

### Temporary and limited CRISPR-Cas adaptation activity in the gut of gnotobiotic mice

The CAPTURE protocol [27] followed by deep sequencing was used again to investigate if the CRISPR array of *E. lenta* DSM 156444 had expanded during colonization in the gut of GB mice. The size of the DNA fragments on an agarose gel suggested expanded CRISPR arrays containing even multiple spacer acquisitions. These expanded CRISPR arrays were observed both in samples from EL+Phage mice (Figure 6), EL+Saline mice, and pure bacterial cultures of the WT *E. lenta* (Figure S8). A protospacer matching the genome of phage PMBT5 as indeed detected in some of the expanded CRISPR arrays (EL+Phage), however, it was only at day 12 and until day 26 that the number of spacers matching the genome of phage PMBT5 were above 100 reads (Figure 6). In contrary to the *in vitro* settings, only one newly acquired spacer (with 75 742 reads out of 76 846 total phage-associated reads, 98.6%) targeted the same phage gene (YP_009807312.1, Figure 6). The PAM for this single protospacer was 5’-TTC (Figure S6). The sequencing yielded a total of 10 716 969 reads of which only 0.7% contained spacer acquisitions that could be assigned to the phage PMBT5 genome. The remaining reads (99.3%) were assigned to PCR products with no spacer acquisition (primer dimers, 89.2%), and to the *E. lenta* genome (10.1%) as also observed in the *in vitro* experiment (Figure S7). Overall, the results indicated a temporary and limited CRISPR-Cas mediated adaptation activity when exposed to phage PMBT5 in a GB mouse model.

**Figure 6:**
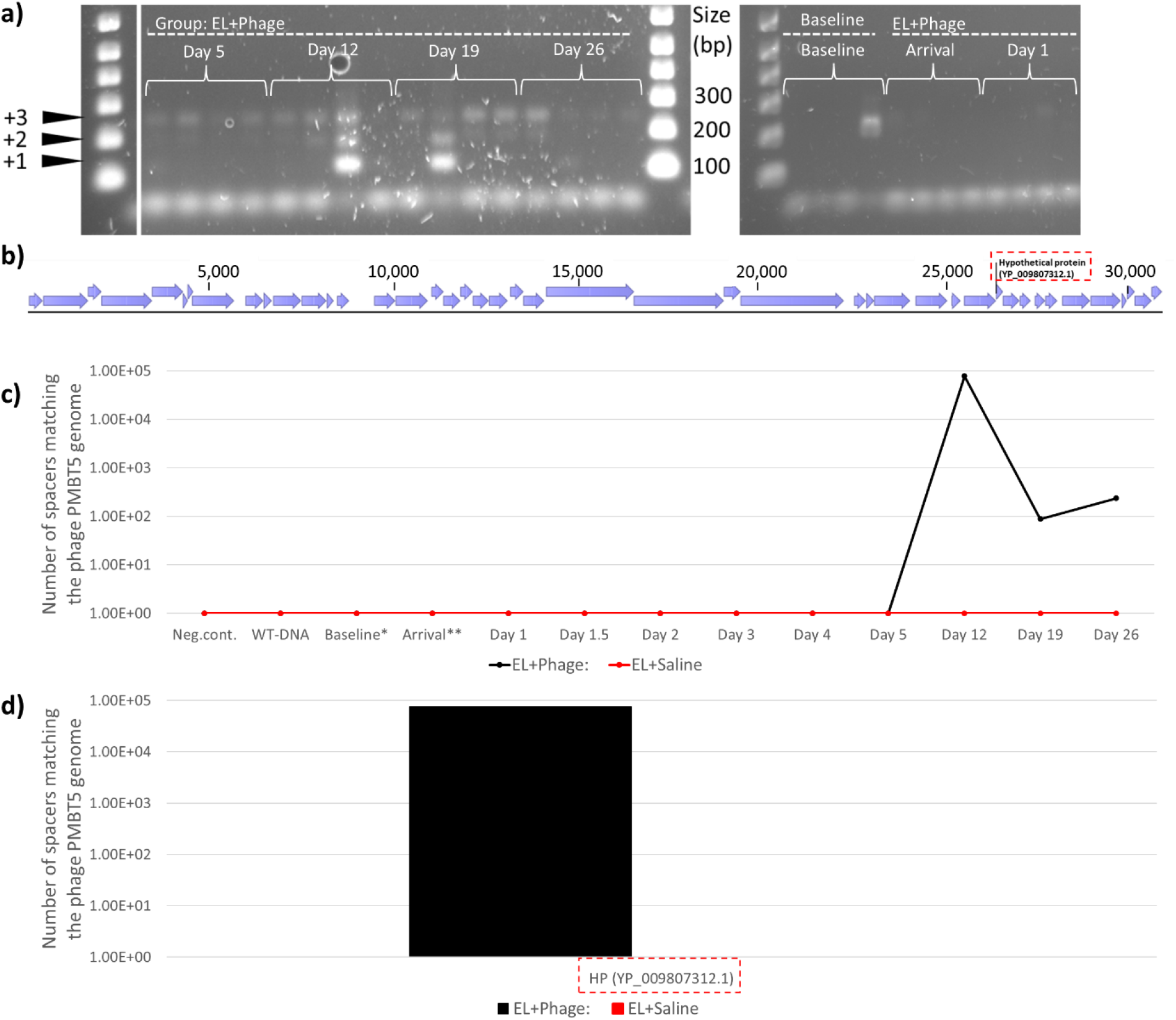
Overview of spacer acquisitions in the *in vivo* settings. a) An agarose gel showing spacer acquisitions in selected samples representing EL+Phage mice from day 5, day 12, day 19, and day 26, as well as from controls at arrival (Day 1) and baseline mice. A 100-bp DNA ladder was used to estimate PCR product size. With the degenerate primers, the acquisition of one spacer “+1” was expected to yield a PCR product at ∼110 bp (Figure S1) and then ∼70 bp for additional spacers. The PCR-product at ∼40 bp likely represented primer dimers. b) The annotated phage genome of PMBT5 highlight the genes that are presented in c) with a line plot showing reads/spacers over time and d) as a bar plot showing the number of reads/spacers that matched at the phage genome. Only one phage gene coding for a hypothetical protein (YP_009807312.1) appeared as a source of spacer*s*. This is marked by a box with red dashed lines. HP = hypothetical protein.

## Discussion

Here we report the activity of a type I-C CRISPR-Cas system harbored by the prevalent human gut bacterium *E. lenta* [41] when exposed to virulent phages in both *in vitro* and *in vivo* settings. With a highly sensitive PCR-based protocol [27] and deep sequencing, we detected MOI-dependent CRISPR-Cas adaptation activity against phage PMBT5 when infecting *E. lenta* DSM 15644 (Figure 3). A decrease in bacterial growth in phage-infected cultures is likely explained by impaired fitness due to acquired phage resistance [42–44] that was not associated to CRISPR-Cas immunity [45]. The bacterial cultures infected at MOIs of 0.1 and 0.01 had a relative higher number of new spacers matching the phage genome as compared to cultures infected at MOIs 10 and 1 (Figure 3). It has been shown that low MOI 0.01 can activate dormancy in archaea and allow them to recover from active CRISPR-Cas immunity [46]. Similar mechanisms of dormancy might explain why *E. lenta* DSM 15644 cultures infected with phage PMBT5 at MOI 0.1 and 0.01 of revealing limited growth, were found to have increased frequencies of spacer acquisitions compared to cultures infected with MOI 10 and 1.

Using *in vitro* settings, 13 protospacers of phage PMBT5 were targeted at all four MOIs of which 3 appeared preferred targets, while only 1 protospacer was targeted in the GB mice. These hotspots of spacer acquisition were within genes coding for a portal protein and three hypothetical proteins (Figures 3 and 6, & Table S7). Based on the 14 protospacers, the adaptation PAM was predicted as 5’-TTC (Figure S6), which is in agreement with another study that predicted similar adaptation PAM for type I-C CRISPR-Cas systems in 15 different *E. lenta* strains using computational approaches [16].

The distinctly different environmental conditions for phage-bacteria interactions in test tubes versus the spatial heterogeneity found in real gastrointestinal conditions in GB mice [47], may explain this clear difference in the number of unique acquired spacers between the *in vitro* and *in vivo* settings. The bacterial to phage ratio in the mouse feces was estimated at 10 (phages were 1 log higher than bacteria, Figure 5), so the decreased frequency of spacer acquisition during *in vivo* conditions may be in line with the low frequency of spacer acquisition observed in the *in vitro* settings with a MOI of 10 (Figures 3 & 6).

While the numbers of reads representing the acquired spacers are arbitrary values due to the basic principles of the CAPTURE protocol [27], it appears that spacer acquisition may be relatively rare in *E. lenta* DSM 15644 when exposed to phage PMBT5 in both *in vitro* and *in vivo* settings. This would also be in accordance with other studies investigating spacer acquisitions [48,49]. Of note, two hypothetical proteins encoded by phage PMBT5 had identical (E-value < 10^−23^) AA domains as four potential anti-CRISPR (acr) protein clusters [38] (YP_009807310.1: cluster 2517 + 20298 and YP_009807284.1: cluster 12618 + 59526) (Figure S9). Therefore, if these phage proteins contain acr features, it might have challenged the detection of spacer acquisitions in *E. lenta* and thereby limited CRISPR-Cas immunity.

In both the *in vitro* and *in vivo* settings, less than 2% of the total sequenced reads could be assigned to the phage PMBT5 genome, while ∼ 25% were assigned to the genome of *E. lenta*, and the remaining reads were primer dimers. The reads that matched to the chromosomal DNA framed almost the entire native CRISPR array (Figure S6) and no other bacterial genes. This phenomenon was detected in all samples independent of the presence of phages. Whether this observation has biological relevance or is just PCR-generated artefacts is not known. Other potential biological explanations may be homologous recombination (driven by the repeats) or a mechanism where the spacers are shuffled to increase spacer diversity at the leader and more expressed end of the CRISPR array. Self-targeting immune memories of CRISPR-Cas have previously been demonstrated [50–52], but does not seem to explain our observation of spacers matching the CRISPR array of the host since no chromosomal genes were targeted.

Using a plasmid system in which we cloned two spacers (S2 and S1) originating from the native CRISPR array of *E. lenta* DSM 15644, we showed clear interference activity of the type I-C CRISPR-Cas system, including the necessity of the PAM 5’-GGG and 5’-TTC (Figure 4). Considering that the adaptation and interference stages consist of different protein complexes being formed, the PAM requirements may be different for both stages [53]. Thereby explaining why both PAM 5’-GGG and 5’-TTC showed high interference efficiency, while 5’-TTC appears to be the preferred PAM that is involved in spacer acquisition. Although, one protospacer was detected with 5’-GGG as PAM (Table S7). The observed interference activity of the type I-C CRISPR-Cas system in *E. lenta* DSM 15644 is in line with another study that reported transcription and interference activity of a type I-C CRISPR-Cas system from the closely related strain *E. lenta* DSM 2243^T^ [16]. Soto-Perez et al. conducted an experimental design using the evolutionary distinct (from *E. lenta*) bacterium *Pseudomonas aeruginosa* and its associated *P. aeruginosa* phages [16]. Whereas we investigated CRISPR immunity of *E. lenta* using natural host-phage relations. Despite the high genomic similarity between *E. lenta* DSM 2243^T^ and DSM 15644 [16], the *E. lenta* DSM 2243^T^ showed no susceptibility against the PMBT5 phage (Figure S10).

The phage PMBT5 was highly virulent *in vitro* since the bacterial culture was completely cleared after phage amplification (Figure S10). It is therefore intriguing, why the bacterial abundance was similar with and without the presence of this seemingly highly virulent phage (Figure 5) during the 26 days in GB mice. Considering that only one new spacer acquisition was detected in GB mice, it suggests that the type I-C CRISPR-Cas in *E. lenta* DSM 15644 does not constitute the main phage resistance strategy in the investigated conditions. Other resistance mechanisms [12,54] are likely involved. In addition, the contribution of physical parameters should not be neglected, since small cavities in the intestinal lumen, mucus production (from host cells) [55,56], protection by numerous bacterial cell layers in microcolony structures [57–59], and the overall spatial distribution in the gut may have protected the bacteria from phages. Avoiding infections would mean less phage resistance, and perhaps even avoiding impaired fitness that is sometimes associated with phage resistance [42–44]. In our mouse model, some sort of equilibrium between bacterial cell division and phage infection of susceptible cells seemed to have occurred (Figure 5). This would be in accordance with a study showing that the spatial heterogeneity of the gut limits predation and favors the coexistence of phages and bacteria [47]. Other studies have also shown host-phage coexistence in other experimental and theoretical settings using the bacterium *Streptococcus thermophilus* and its lytic phage 2972 [60,61].

Although, the CRISPR-Cas system only provided limited phage immunity, this is the first study to show the activity of the type I-C CRISPR-Cas system in *E. lenta* targeting its antagonist phage in both *in vitro* and *in vivo* settings.

## Acknowledgements

We thank Professor Alejandro Reyes (Universidad de los Andes, Colombia) for initial discussions of the design of the gnotobiotic mouse (GB) model and we also thank Dr. Frank Hille (Max Rubner-Institut, Germany) for suggestions that improved the initial draft of the manuscript. In addition, we are grateful to Helene Farlov and Mette Nelander at Section of Experimental Animal Models (University of Copenhagen, Denmark) for handling GB mice in the isolator, as well as the staff at Department of Experimental Medicine (University of Copenhagen, Denmark) for their cooperation in housing the GB mice. We also thank lab technicians Pernille Lærke Jensen and Emilie Gaal at Section of Microbiology and Fermentations (University of Copenhagen, Denmark) for helping with the interference activity assay.

## Funding

Funding was provided by the Danish Council for Independent Research with grant ID: DFF-6111- 00316 (PhageGut). This work is supported by the Joint Programming Initiative ‘Healthy Diet for a Healthy Life’, specifically here, the Danish Agency for Science and Higher Education. S.M. acknowledges funding from the Canadian Institutes of Health Research (Team grant on Intestinal Microbiomics, Institute of Nutrition, Metabolism, and Diabetes). S.M. holds a Tier 1 Canada Research Chair in Bacteriophages.

## Author contributions

TSR, DSN, AKH, and SM conceived the research idea and designed the study; TSR and AKK performed the experiments; TSR, AKK, DSN, SM, GR, WK, LD, MKM, SS, HN, CMAPF, and FKV performed laboratory and data analysis; TSR wrote the first draft of the manuscript. All authors critically revised and approved the final version of the manuscript.

## Competing interests

All authors declare no conflicts of interest.

## Supplemental materials

All supplemental materials (figures, tables, primer lists, additional methods etc.) are available through the link https://osf.io/n5dqj/?view_only=bf883f5b3c2444d58acd7409d797d9a0.

## Notes

https://osf.io/n5dqj/?view_only=bf883f5b3c2444d58acd7409d797d9a0

### Competing Interest Statement

The authors have declared no competing interest.

