## Supplemental figures for "CRISPR-Cas provides limited phage immunity to a prevalent gut bacterium in gnotobiotic mice"

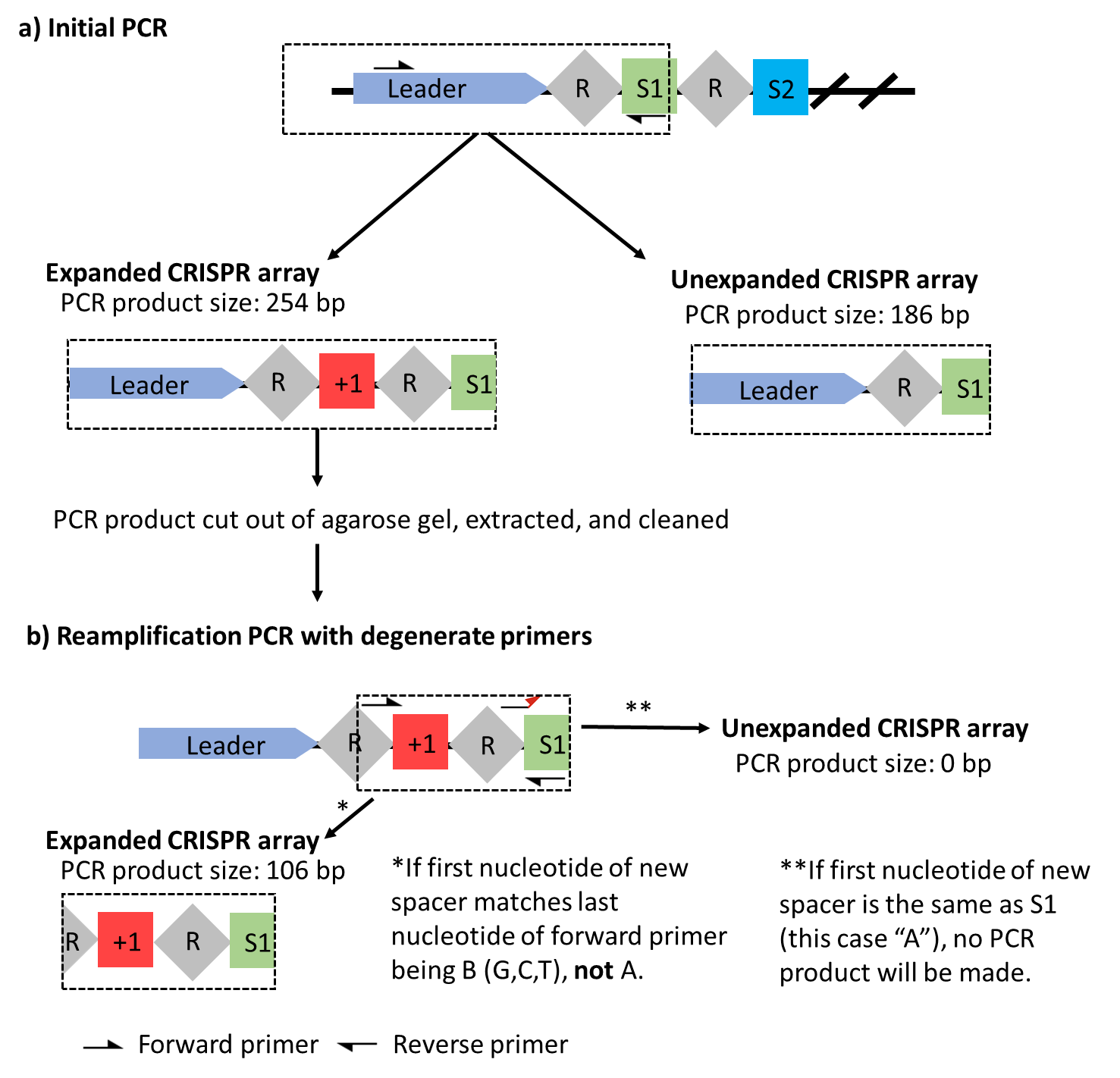

Figure S1: A graphical representation of the reamplification PCR procedure (CAPTURE) with degenerate primers used to detect expanded CRISPR arrays. a) Initial PCR: Primers are matching parts of the leader and spacer 1 (S1), and in the case of expanded CRISPR array the PCR product size will be 254 bp. Without being able to see the “expanded CRISPR array”, a piece of the gel is cut out, DNA extracted and cleaned. b) The cleaned PCR-product is then subjected to reamplification with degenerate primers, but these primers are binding to the CRISPR array repeat with a nucleotide overhang of G, C, or T, and a S1 primer as in the initial PCR. If the CRISPR array is expanded and if the new spacer sequence starts with G, C, or T, the degenerate primers will bind and produce a PCR product with a size of ~106 bp.

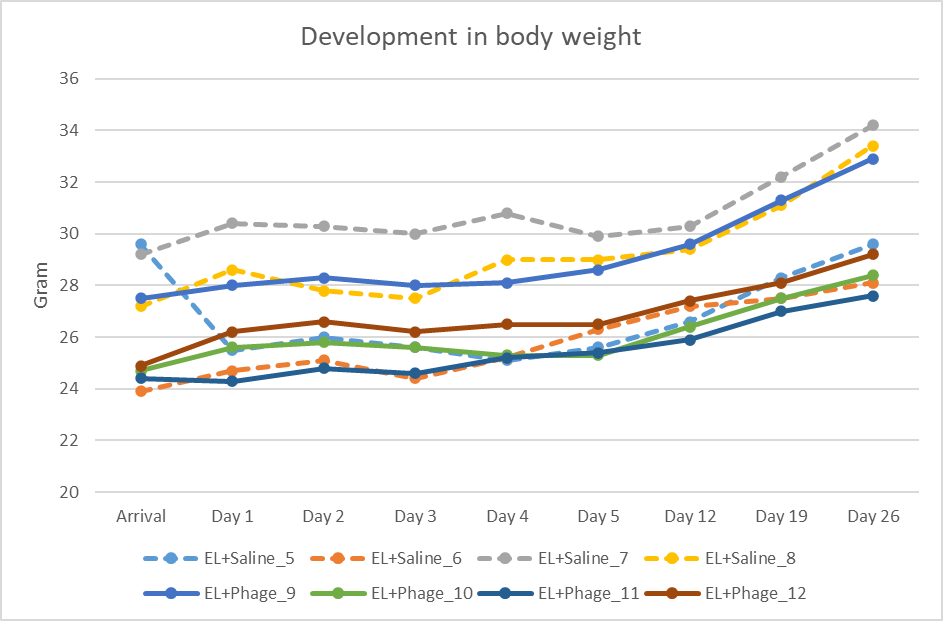

Figure S2: A line plot showing the development of the body weight in grams of the individual gnotobiotic mouse (numbered from ID 5-12) that was monitored throughout the study.

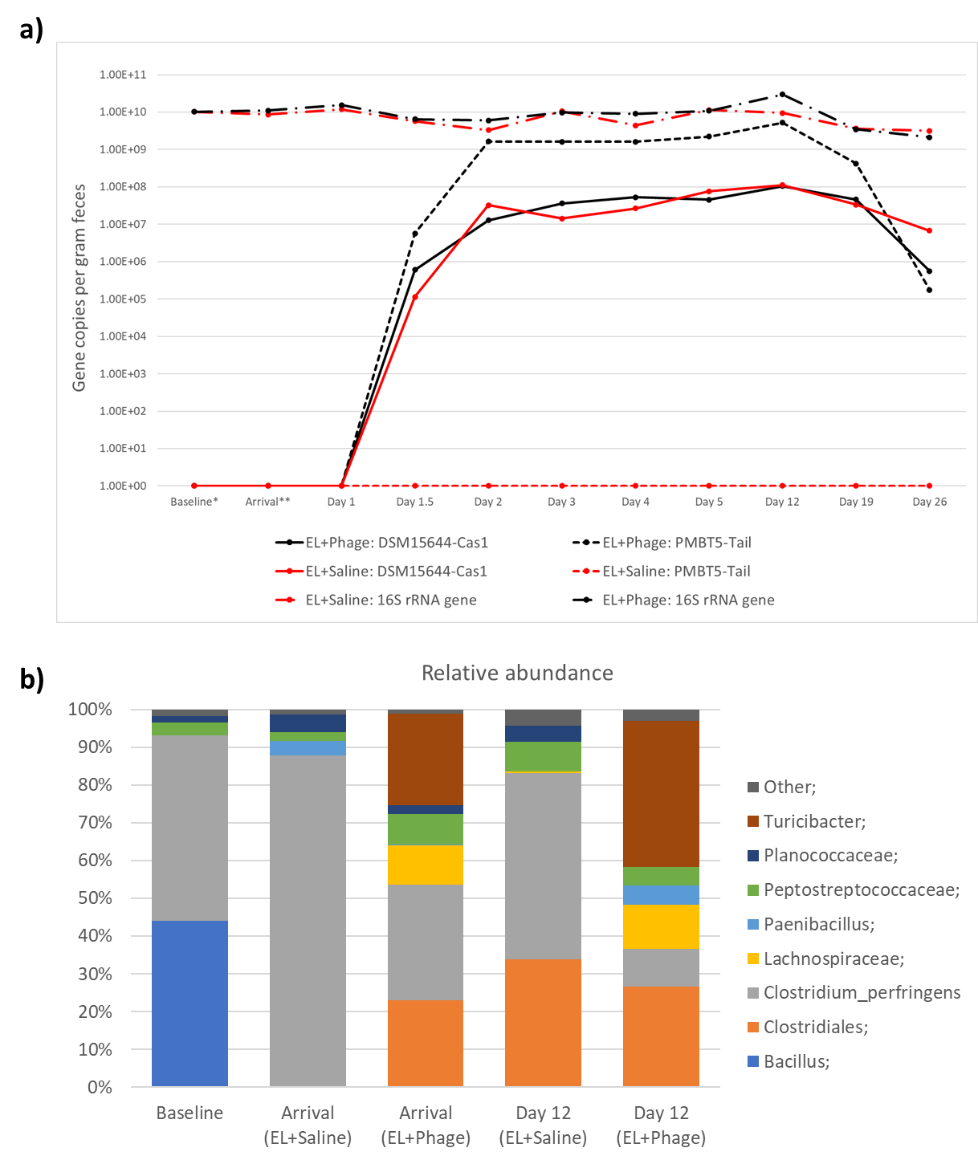

Figure S3: qPCR analysis and full 16S rRNA gene sequencing. a) Bacterial and phage abundance in feces samples at different time points measured by qPCR. Primers were designed to specifically target the genomes of *E. lenta* DSM 15644 and phage PMBT5. Universal 16S rRNA gene primers (V3 region) were used to measure the total 16S rRNA gene copies found in the feces samples. A threshold of minimum 10 gene copies was applied. Surprisingly, the qPCR showed more than 10^9^ 16S rRNA gene copies per gram feces in the GB mice before inoculation with E. lenta DSM 15644. b) Relative abundance (in percentage) of bacterial taxa found in the GB mice at three selected time points: Baseline (Feces samples from GB mice euthanized at the age of 3 weeks), Arrival (feces samples from GB mice when transferred from isolator to individual ventilated cages at another housing facility), and Day 12 after inoculation. The full 16S rRNA gene sequencing revealed bacterial taxa such as *Paenibacillus* (spore forming)*, Clostridium perfringens* (spore forming)*,* and *Bacillus* (spore forming). We speculate that these microbes represent taxa that were killed during sterilization of the feed but still detected, due to the lack of live microbes in the gut of the GB mice.

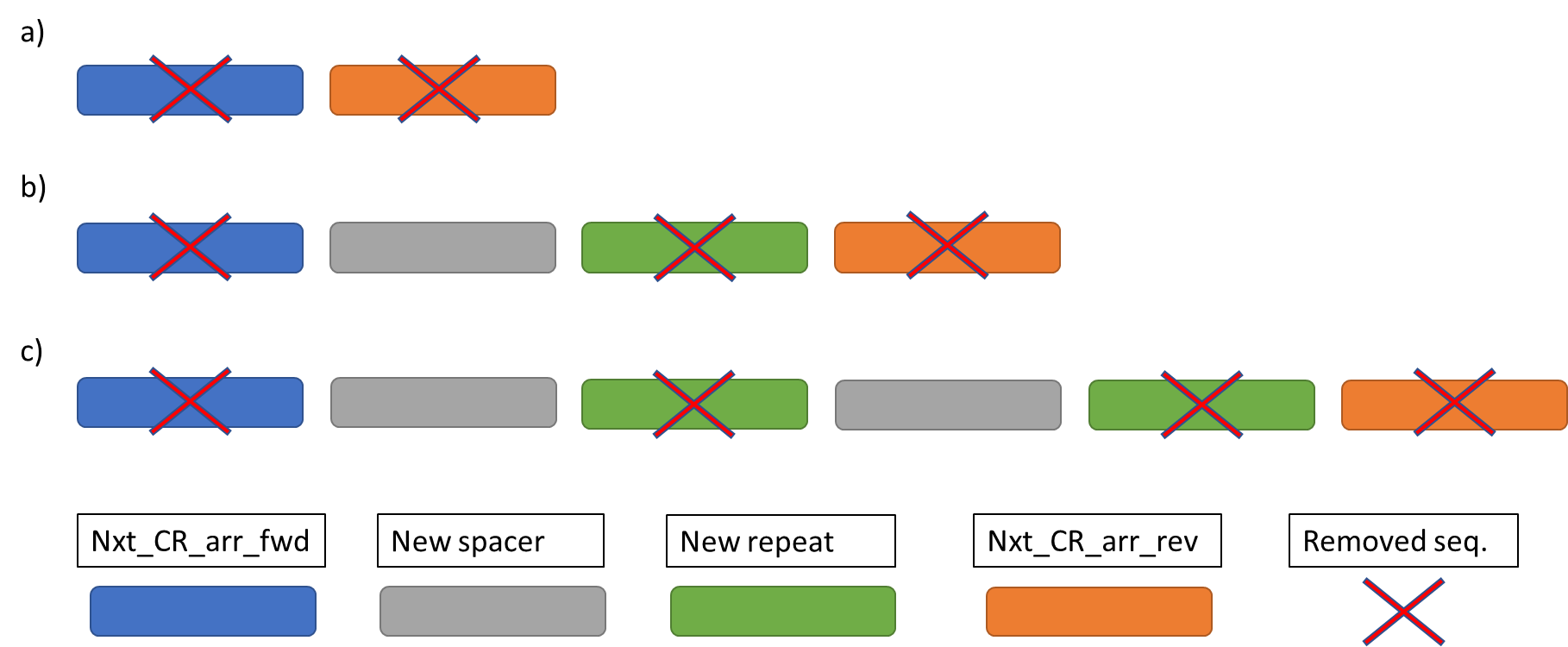

Figure S4: Visualization of how redundant sequences (adaptor and repeats) were removed from the raw Illumina NextSeq reads. a) no spacer acquisition, b) one spacer acquisition, c) two spacer acquisitions, etc.

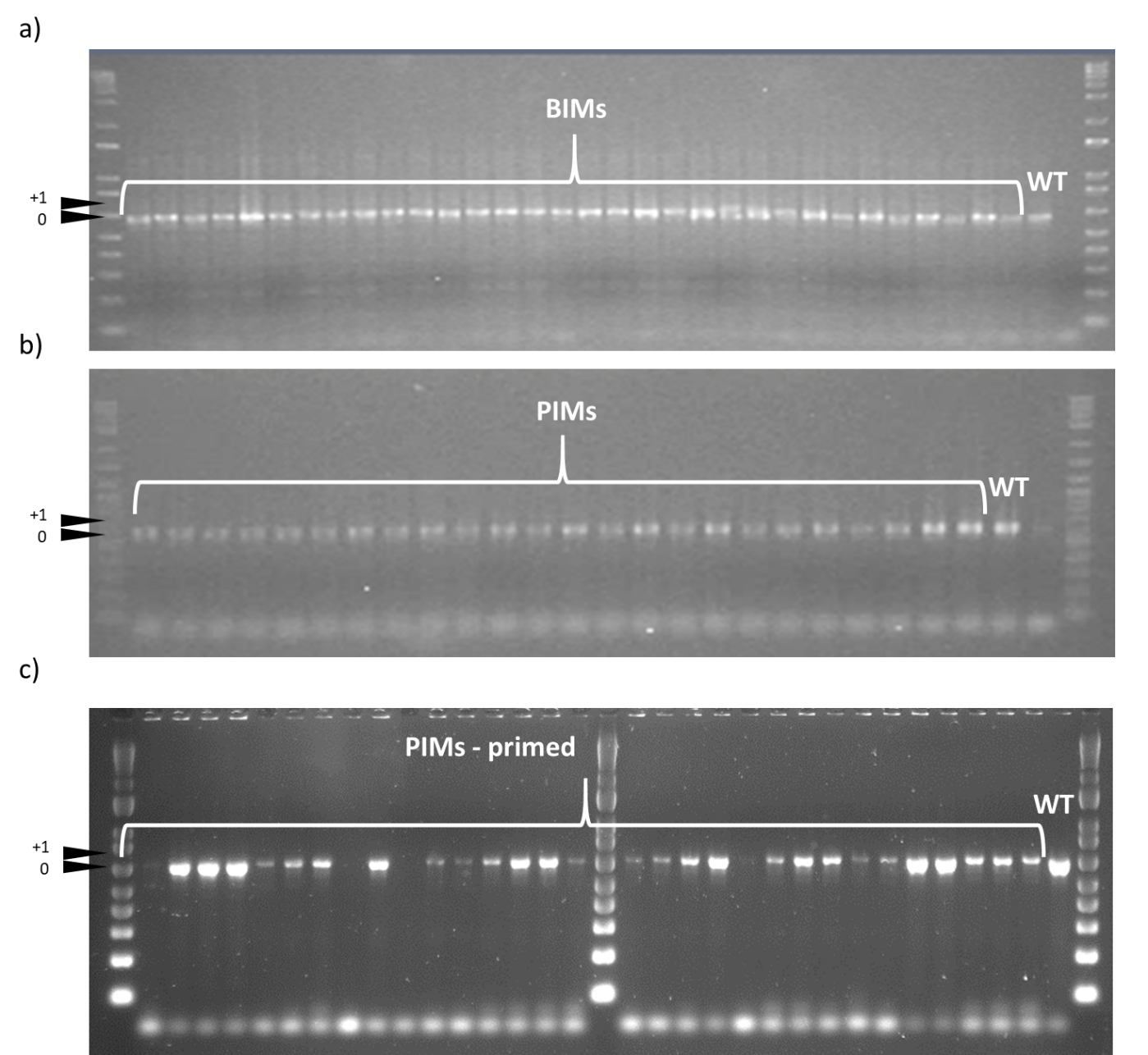

Figure S5: Results of selected gels used for detection of spacer acquisitions at the 5’-end of the CRISPR array (primer set: CRISPR_ac_5). a) BIM assay, b) PIM assay, and c) primed adaptation. No spacer acquisition was detected by PCR in any of the assays. The black arrows mark the PCR-product size of either no spacer acquisition (0) or one spacer acquisition (+1). WT = wild-type, *E. lenta* DSM 15644. The reference DNA ladder is a 100-bp scale.

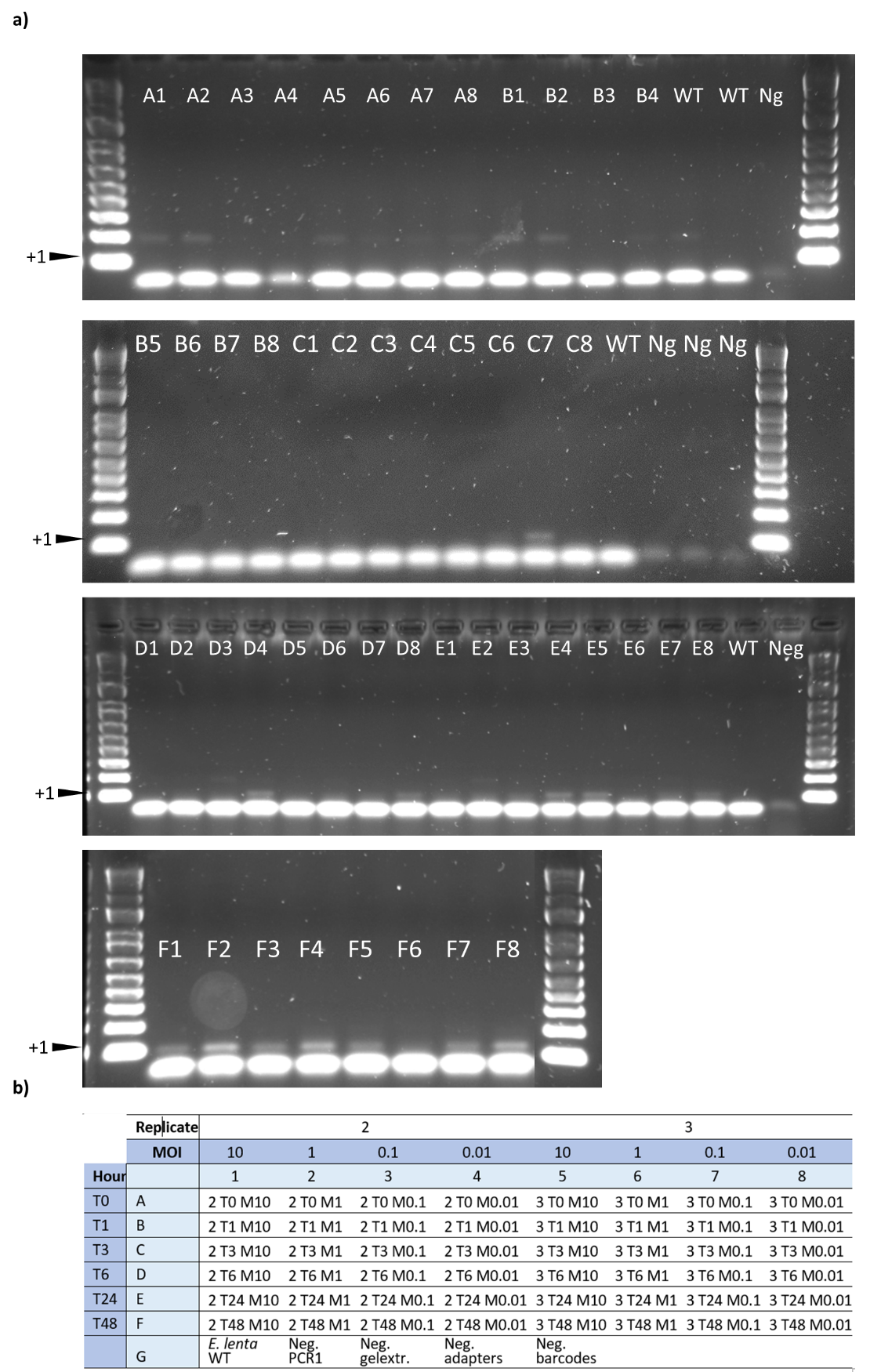

Figure S6: Detection of PCR products of all individual samples. a) A DNA band at +1 arrow marks PCR-products containing one spacer acquisition. The reference DNA ladder is a 100-bp scale. b) Table lists sample information of replicate number, time of sampling, and MOI of each sample on the gel images. WT = Wild-type. Neg/Ng = negative control.

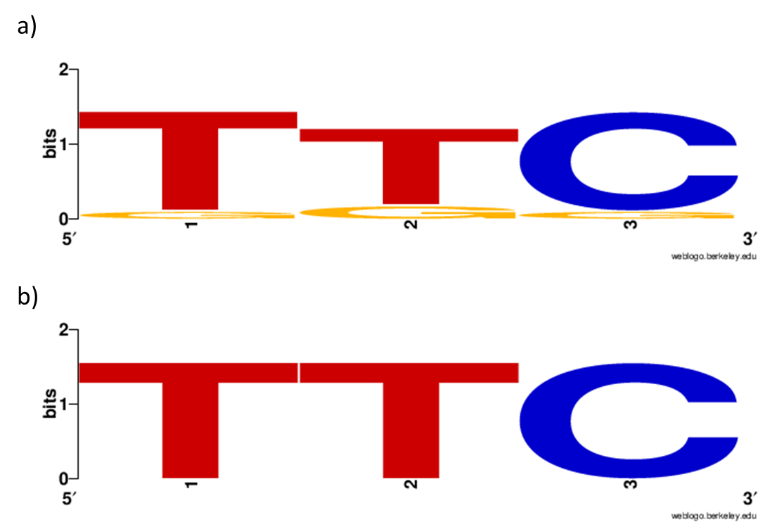

Figure S7: Sequence logo of a) the three-nucleotide predicted PAM associated to the 13 protospacers detected on phage PMBT5 genome in the *in vitro* experiment and b) of the only protospacer observed for the *in vivo* experiment.

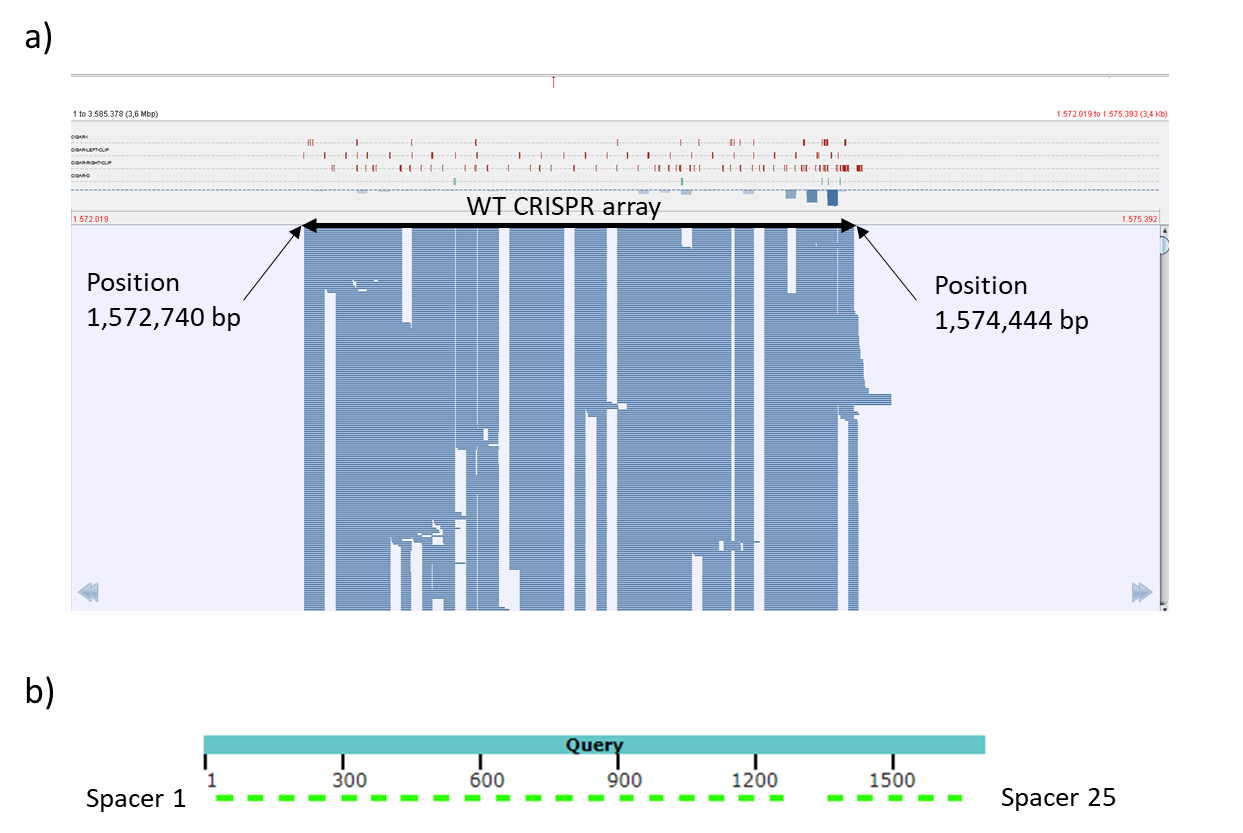

Figure S8: Alignment of reads/spacers against the genome of *E. lenta* DSM 15644. a) Alignment of reads/spacers that matched to the chromosomal DNA specifically framing the native CRISPR array (position 1,572,740 - 1,574,444 bp). This event was observed at all four MOIs as well as the control without phages in the *in vitro* settings. Similar phenomenon was also observed in the *in vivo* settings. With the available data it is not possible to document whether this observation is associated to a biological relevance or represent a PCR generated artefact. b) BlastN alignment showing that 24 out of the 25 native spacers matched reads/spacers with an identity of 100%.

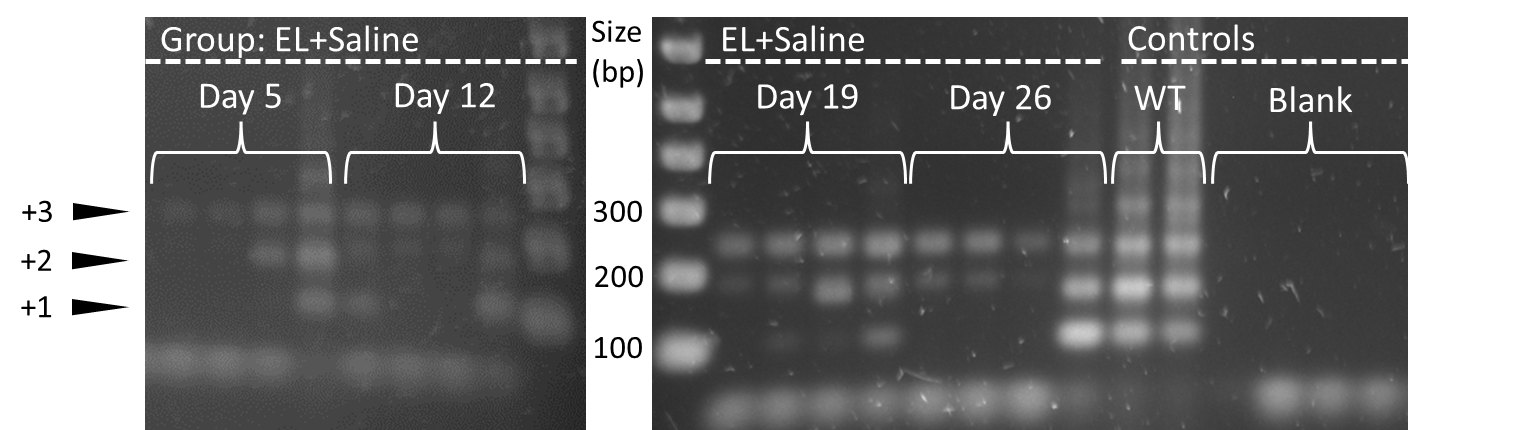

Figure S9: An agarose gel showing spacer acquisitions in selected samples of EL+Saline mice from day 5, day 12, day 19, and day 26. Controls were the WT *E. lenta* DSM 15644 and blank samples. DNA ladder is 100-bp scale. With the degenerate primers, the acquisition of one spacer “+1” was expected to yield a PCR product at ~110 bp (see Figure S1) and then ~70 bp for additional spacers. The PCR-product at ~40 bp likely represented primer dimers.

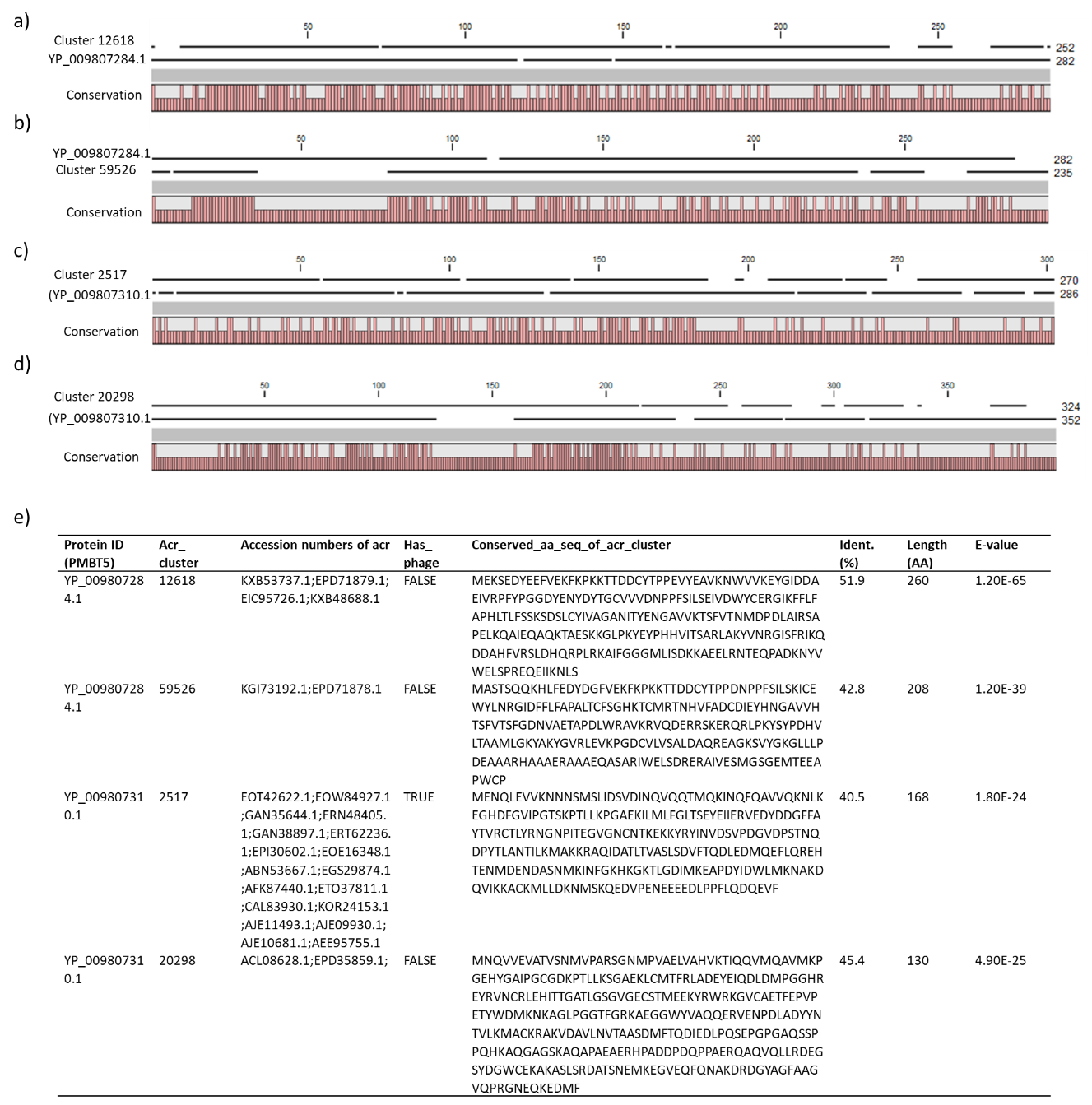

Figure S10: Protein sequences from phage PMBT5 with shared domains and conserved amino acids (AA) to putative anti-CRISPR (acr) clusters. a) YP_009807284.1 and acr cluster 12618, b) YP_009807284.1 and acr cluster 59526, c) YP_009807310.1 and acr cluster 2517, d) + YP_009807310.1 and acr cluster 20298. e) A table listing the accession ID of the PMBT5 and acr protein sequences along with the identity percentage, alignment length and E-value. According to Gussow et al. 2020: TRUE = acr cluster that is associated to a characterized phage, FALSE = acr cluster only based on computational prediction.

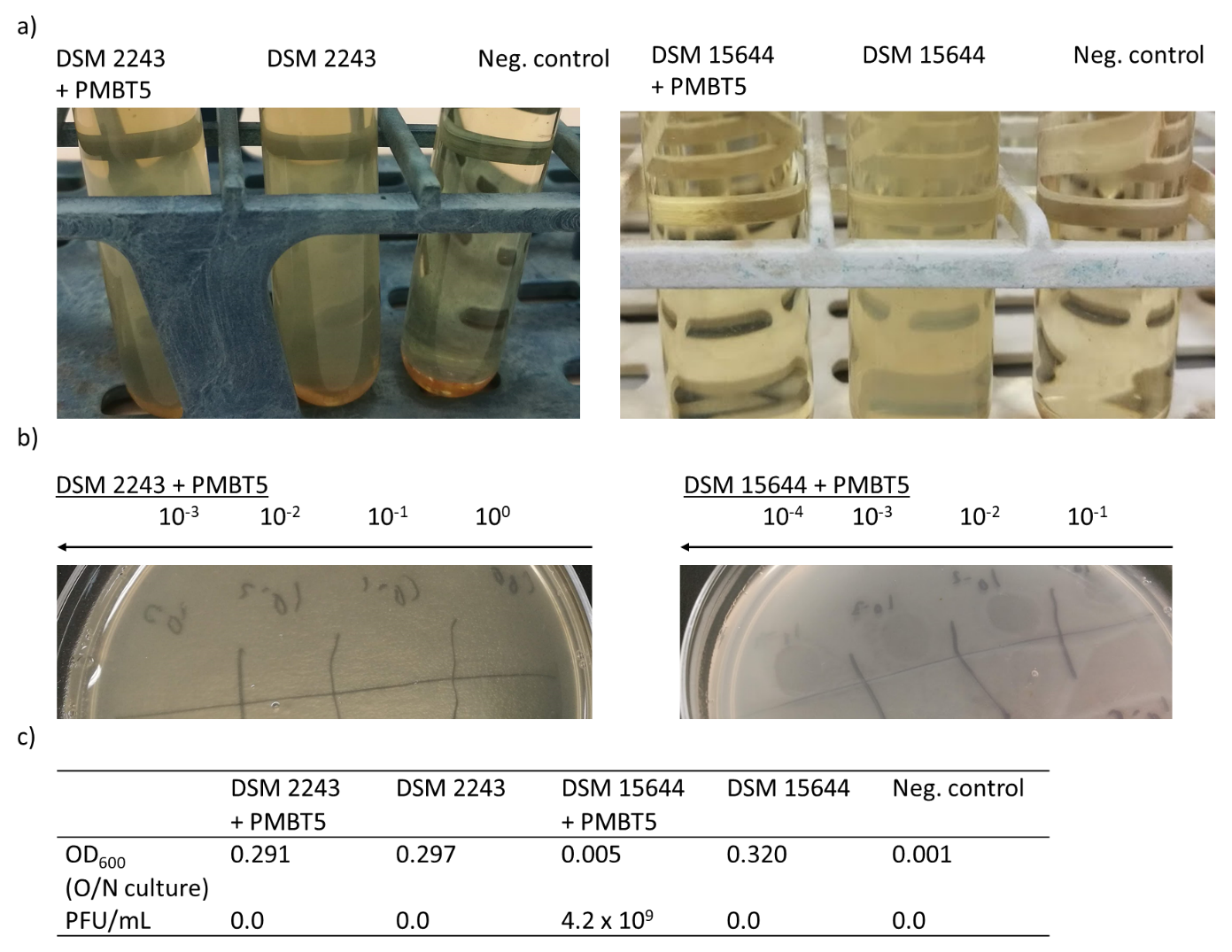

Figure S11: Phage PMBT5 infectivity test by measuring the culture optical density a) and titers (PFU/mL) on *E. lenta* DSM 15644 and *E. lenta* DSM 2243 b). Phage propagation in liquid was at an initial MOI of 1. c) Average OD_600nm_ and PFU/mL values of triplicates.
