## Supplemental methods for "CRISPR-Cas provides limited phage immunity to a prevalent gut bacterium in gnotobiotic mice"

### Assays for CRISPR-Cas spacer naive spacer acquisition activity

The investigation of natural adaptation through spacer acquisition by bacteriophage-insensitive mutants (BIMs) were conducted as previously described [1]. CRISPR activities can be prevented by anti-CRISPR (acr) proteins harboured by phages [2], however this can be circumvented by introducing a plasmid containing both a marker gene (e.g. antibiotic resistance gene) and spacer sequences originating from the bacterial CRISPR array into the bacteria. Introduction of such plasmids can generate in some cases plasmid interfering mutants (PIMs) [3] that have acquired a new spacer originating from the plasmid. Plasmid pNZ123 containing a chloramphenicol resistance gene as a marker [4,5] was introduced into *E. lenta* DSM 15644 by electroporation [6] and PIM assays were performed as described previously [3]. Primed adaptation has also been shown in other type I-C CRISPR-Cas systems [7], thus three DNA variants of spacer 1 (S1) with a recognized PAM (5’-GGG) (Table S1) were designed to induce priming [8]. pNZ123 constructs were also transformed into *E. lenta* DSM 15644 by electroporation [6]. Selection of colonies with potential primed adaptation were selected as PIMs [3], with chloramphenicol selection marker. Screening of natural or primed adaptation of new CRISPR spacers in selected colonies was performed by PCR as described earlier [1,3] using DreamTaq Green PCR (Thermo Scientific), with three primers (Table S1) designed to screen for spacer acquisitions at the 5’-end (CRISPR_ac_5), 3’-end (CRISPR_ac_3), and the middle (CRISPR_ac_mid) of the CRISPR array since the spacer acquisition behaviour was not yet known in this bacterial species.
